# Activation of stably silenced genes by recruitment of a synthetic de-methylating module

**DOI:** 10.1101/2021.09.11.459888

**Authors:** Wing Fuk Chan, Christine R. Keenan, Timothy M. Johanson, Rhys S. Allan

**Author notes:** Correspondence: Rhys S Allan and Wing Fuk Chan., Immunology Division, Walter and Eliza Hall Institute of Medical Research, 1G Royal Parade, Parkville, Victoria, Australia 3052.

## Abstract

Stably silenced genes that display a high level of CpG dinucleotide methylation are refractory to the current generation of dCas9-based activation systems. To counter this, we created an improved activation system by coupling the catalytic domain of DNA demethylating enzyme TET1 with transcriptional activators (TETact). TETact induces transcription of heavily suppressed non-coding RNA and surface protein, and the reactivation of embryonic haemoglobin genes in non-erythroid cells.

---

Clustered regularly interspaced short palindromic repeats and the associated Cas9 endonuclease (CRISPR/Cas9) represent a transformative and programmable tool to modify the genome^1^. Through Watson-Crick base pairing, the RNA-guided Cas9 can target the genome ubiquitously, as long as a very short protospacer adjacent motif (PAM) is present. Cas9 was further engineered to remove nucleolytic activity (dCas9) and repurposed as a DNA-binding platform^1-3^. As such, gene transcription can be induced by recruiting transcriptional activators to dCas9 via direct fusion or indirect tethering. While fusion of a single activation domain VP64 causes only modest gene upregulation^4, 5^, the second generation CRISPR activators involve recruitment of multiple effectors, of which the dCas9-VPR^6^, SunTag-VP64^7^ and synergistic activation mediator^8^ (SAM) appear to be the most potent systems^4^.

Programmable gene activation has led to a plethora of applications, including dissection of gene function^1, 3, 9^, genetic screening for important coding or non-coding elements^1, 3, 9^, programmed cellular differentiation^6^ and curative therapeutics^1, 3, 9^. Such applications require the robust activation of candidate genes regardless of the repressive elements present at the relevant loci, including DNA methylation^10^. Thus, any system that can expand our ability to remove or circumvent these repressive elements has obvious value.

In a previous study, we characterised a long non-coding RNA species *Dreg1* within the enhancer region of *Gata3*^11^. Expression of *Dreg1* is highly correlated with *Gata3* expression being expressed in T cell subsets, but completely and stably silenced in B cells. To gain insight into *Dreg1* function, we attempted to activate it in a murine B cell line (A20) using second-generation CRISPR activation systems, SAM and SunTag-VP64. Unfortunately, targeting the *Dreg1* TSS with either SAM or SunTag-VP64 failed to activate transcription (Fig. 1a).

**Figure 1.**
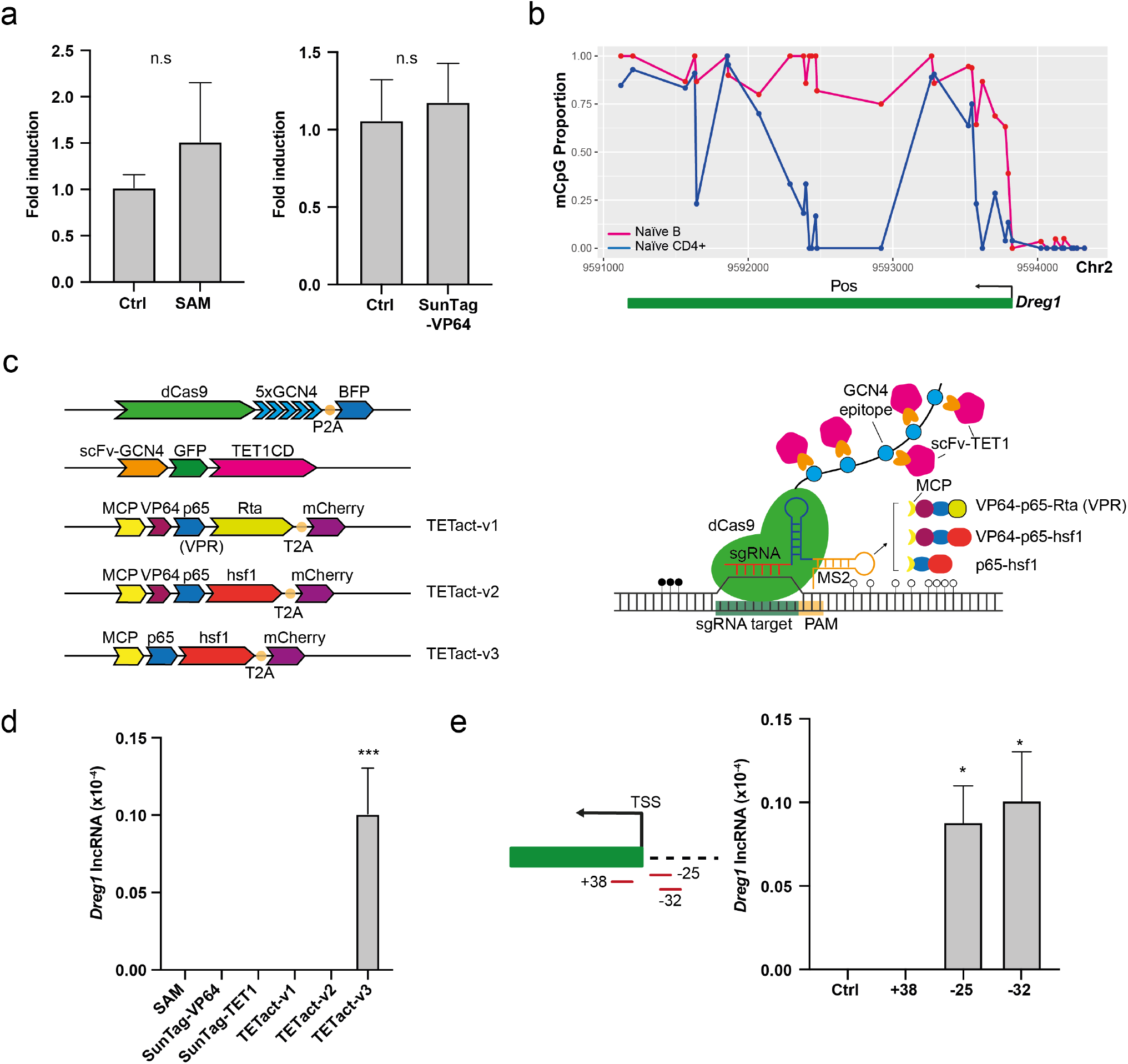
Activation of the T-cell specific lncRNA *Dreg1* in A20 B cells. (**a**) Fold activation of *Dreg1* in A20 cells transduced with sgRNA targeting *Dreg1* promoter together with SAM or SunTag-VP64 constructs. Fold change is calculated by ΔΔCt method. (**b**) DNA methylation (mCpG) profiles of naïve B and CD4+ T cells at the *Dreg1* locus, plotted as population proportion of methylated cytosine in each CpG dinucleotide motif. (**c**) Schematics of TETact systems and corresponding construct designs – multiple copies of TET1CD are recruited to dCas9 via the GNC4 epitopes, whereas the activator domains (v1 – VPR, v2 – VP64-p65-hsf1, v3 – p65-hsf1) are recruited via 2 MS2 aptamers. (**d**) *Dreg1* lncRNA expression in A20 cells transduced with sgRNA targeting *Dreg1* promoter in different systems as indicated. (**e**) Activation of *Dreg1* lncRNA using different sgRNA targeting location. Expression level is relative to β-actin (*Actb*) level as 2^-ΔCt^. Data shown are mean ± s.e.m. from 3 independent transductions. n.s., non-significant, *P < 0.05, ***P < 0.001

Interestingly, activation of other lncRNAs using the second-generation CRISPR activators only leads to very low or modest upregulation^4, 8^ and we postulated that DNA methylation may be an impediment to efficient activation of these genes^12^. The DNA methylation pattern of the *Dreg1* locus in T and B cells was determined via publicly available whole genome bisulphite sequencing (WGBS) data^13^. As predicted, regions around the *Dreg1* TSS and gene body are differentially methylated (Fig. 1b) between the two cell types, with most CpG dinucleotides in B cells being heavily methylated.

This prompted us to investigate the possibility of activating a heavily methylated and repressed *Dreg1* by simultaneously recruiting the DNA demethylating enzyme TET1^14^ and transcription activators to the target site. A recent study utilised a direct fusion of the catalytic domain of TET1 (TET1CD) to dCas9 to reactivate synthetically silenced genes^15^. However, due to the large size of TET1CD, direct fusion to dCas9 together with a selection marker is unfavourable in the context of immune cells or for therapeutic application, as it likely exceeds the cargo limit of lentiviral vectors. In addition, previous studies have suggested that more efficient gene activation is achieved with multiple copies of TET1CD^16^. We therefore adopted the previously described SunTag approach for the recruitment of TET1CD^16^, and the RNA aptamer MS2 harboured within the sgRNA for the recruitment of different combinations of transcription activators herein designated as TETact (Fig. 1c, TETact v1-v3). Of the three combinations tested, the fusion of MS2 coat protein with the bipartite activator (p65-hsf1, v3) are most effective in inducing *Dreg1* transcription, from an undetectable level in A20 controls, expression was significantly upregulated to 1/100000 of β-actin level (Fig. 1d). Surprisingly, recruitment of tripartite activators (VP64-p65-hsf1 or VPR) failed to activate the lncRNA, possibly due to steric hindrance imposed by the larger size of these tripartite activators.

Next, we tested the effects of module position on activation strength by designing sgRNAs targeting 3 different sites around the TSS (Fig. 1e). As predicted, activation is extremely sensitive to the target site location in relation to the TSS. While sgRNAs located upstream of the TSS robustly induced *Dreg1* expression, activation did not occur when the sgRNA target site was towards downstream of the TSS (Fig. 1e). Given that the TSS of many lncRNAs and enhancer RNAs are poorly annotated, these experiments suggest that caution, as mistargeting by only 10s of base pairs can cause failure of activation.

To further characterise TETact, we next performed a detailed assessment of the efficiency and kinetics of activation of CD4, a surface protein that defines a subset of T cells.. Again utilising WGBS data^13^, a differential DNA methylation pattern was observed at the CD4 promoter between B and T cells (Supplementary Fig. 1), with the highly methylated DNA in B cells consistent with the lack of transcription of CD4 in this cell-type. We designed sgRNAs targeting the CD4 promoter in A20 cells using different CRISPR activation systems and monitored expression by flow cytometry for 14 days. As predicted, second-generation activators failed to drive a high level of CD4 expression (Fig. 2b, c and Supplementary Fig. 2). As such, only 10% of SAM population (P<0.01 vs control, *t*-test) showed detectable surface CD4 expression (MFI ∼ 500). Similarly, SunTag-VP64 cells showed minimal detectable surface CD4. In stark contrast, approximately 80% of the cells containing TETact with the bipartite activator (v3) exhibited surface CD4 (MFI ∼ 20000) from day 4 (P<0.001, *t*-test)(Fig. 2a-c). In this population significant activation was seen as early as day 2 post sgRNA transduction, with 50% of cells exhibiting detectable surface CD4 (P<0.01, *t*-test). Bisulphite sequencing of the CD4 promoter in the A20-TETact-v3 cells confirmed the engineered demethylation of the region (Fig. 2d). On the other hand, when tripartite activators (TETact-v1 & v2) were recruited, activation was less effective, with these cells showing a lower percentage of CD4+ cells with a lower expression level (Fig. 2b, c and Supplementary Fig. 3). Of note, by recruiting TET1CD alone (SunTag-TET1), CD4 expression became detectable on d7 to d14 (P<0.001, *t*-test), suggesting DNA methylation indeed plays a role in suppressing CD4 in B cells (Fig. 2b, c and Supplementary Fig. 3).

**Figure 2.**
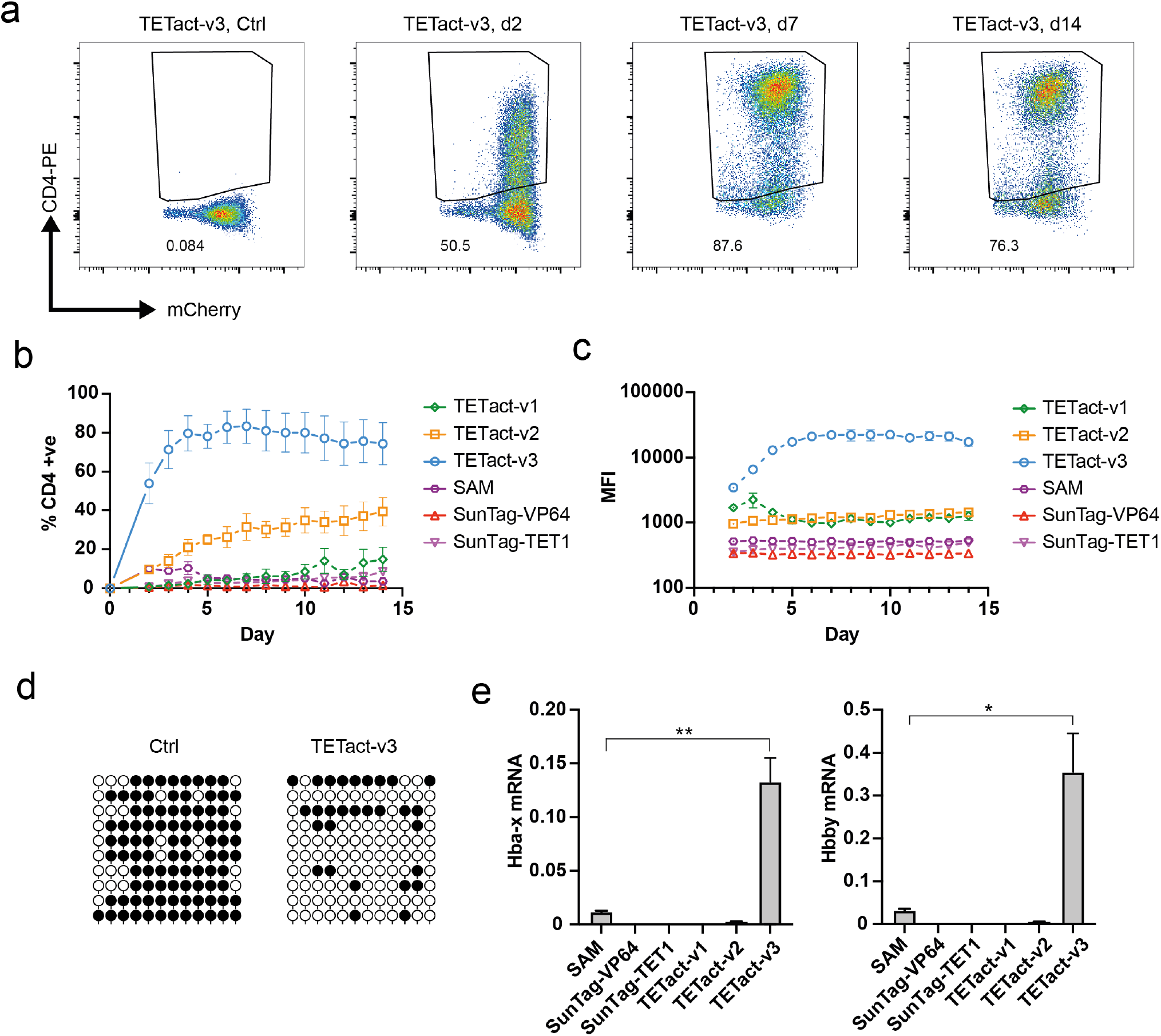
TETact is a potent activator of stably silenced genes. (**a**) Representative flow cytometry plots showing CD4 surface expression in A20-TETact-v3 cells transduced with CD4-targeting sgRNA on the indicated day post-sgRNA-transduction. (**b**) Percentage of population with surface CD4 expression over a 14-day time course for A20 cells with various activators as indicated. (**c**) Median fluorescence intensity (MFI) of CD4-PE over a 14-day time course. (**d**) Bisulphite sequencing of CD4 promoter for A20-TETact-v3 cells transduced with either control or CD4-targeting sgRNA. Open lollipops indicate nonmethylated CpG dinucleotides whereas closed lollipops represent methylated dinucleotides. Each row represents an analysed clone. Ten clones were analysed in each group. (**e**) Expression of *Hba-x* and *Hbb-y* in A20 cells transduced with the promoter-targeting sgRNA in different systems as indicated. Expression level is relative to *Actb* as 2^-ΔCt^. Data are shown as mean ± s.e.m. from 3 independent transductions. *P < 0.05, **P < 0.01

To further explore the potential application of TETact, we attempted to activate embryonic globin genes in ‘adult’ cells, a major aim of gene therapy to treat hemoglobinopathies^17-19^. Adult haemoglobin is composed of α and β chains and mutations in these genes can lead to various blood disorders, for instance, α- and β-thalassaemia as well as sickle cell anaemia^20^. In contrast, during embryonic development haemoglobin is instead composed of other globin chains and reactivation of these represents a promising therapeutic cure for such disorders^17-19^. WGBS data showed highly methylated DNA across both loci in both B and T cells (Supplementary Fig. 4). We therefore designed sgRNAs to target the murine embryonic α-like ζ-globin (*Hba-x*) and β-like ε_y_-globin (*Hbb-y*) in the A20 cell line. qRT-PCR analysis revealed that TETact-v3 outperformed the other systems in upregulating *Hba-x* and *Hbb-y* (Fig. 2e).

The existing second generation CRISPR activators induce transcription through recruitment of various chromatin modifying proteins and transcription factors which alter the local epigenetic landscape such as histone modifications and nucleosome spacing^6, 8^. However, we found that these systems were inefficient at activating genes that contained high levels of methylated CpG dinucleotides. Here we demonstrated that simultaneous recruitment of DNA demethylating enzymes and activation domains can lead to a more robust transcriptional activation of stably silenced genes. Coincidentally, a similar system CRISPRon was developed early this year with direct fusion of a single TET1CD to dCas9 and recruitment of VPR through an RNA scaffold^15^. Whilst this system was able to reverse the repressive state rendered by CRISPR-mediated stable silencing, it is yet to be demonstrated to be able to activate stably and naturally silenced genes. We suspect that the multiple copies of TET1CD recruited by the SunTag epitope in our TETact system are required for robust and efficient gene activation in these settings. The utilisation of SunTag also enables gene delivery via lentivirus, which would be more favourable in certain biological contexts.

Importantly, the activation of stably silenced genes has many important applications from studies of fundamental biology through to gene-editing therapeutics and cellular reprogramming. The robust gene activation from the TETact system presented here will facilitate these applications.

## Online Methods

### Cell culture

The A20 cell-line was cultured in RPMI 1640 with 2 mM GlutaMAX, 50 µM β-mercaptoethanol and 10% heat-inactivated foetal calf serum (FCS). HEK 293T cells were cultured in DMEM with 2 mM GlutaMAX and 10% heat-inactivated FCS without antibiotics.

### Plasmid design and construction

The lentiviral vector dCas9-5xGCN4-BFP was constructed by amplifying the GCN4 array from pCAG-dCas9-5xPlat2AfID (Addgene #82560) with primers bearing the BamHI and NotI sites at the 5’ and 3’ end respectively, and cloning into the corresponding site in pHRdSV40-dCas9-10xGCN4-P2A-BFP (Addgene #60903). Plasmid scFv-GCN4-sfGFP-TET1CD was constructed by cloning sfGFP-TET1CD fragments from pCAG-scFvGCN4sfGFPTET1CD (Addgene #82561) with BamHI and NotI cuts to the corresponding sites in pHRdSV40-scFv-GCN4-sfGFP-VP64-GB1-NLS (#60904). MCP-p65-hsf1-mCherry was constructed from Addgene plasmid MS2-P65-HSF1_GFP (#61423) by replacing GFP with an mCherry gene. Vector gRNA-MS2×2-TagRFP657 was constructed from pLH-sgRNA1-2XMS2 (Addgene #75389) by removing the ccdB and replacing with a shorter BbsI cloning cassette, made from annealing complementary oligos, to the BbsI site in the plasmid, hygromycin resistance gene was further replaced with a TagRFP657 gene obtained from pMSCVpuro-TagRFP657 (Addgene #96939). Based on MCP-p65-hsf1-mCherry, plasmids MCP-VP64-p65-hsf1-mCherry and MCP-VPR-mCherry were constructed via In-Fusion Cloning with VP64 or VPR obtained from the Addgene plasmid #84244.

For SAM activation, vector dCas9-VP64-mCherry was modified from Addgene plasmid dCas9-VP64-GFP (#61422) by exploiting NheI and EcoRI sites to replace the GFP with an mCherry gene. MCP-p65-hsf1-BFP was modified from Addgene plasmid MS2-P65-HSF1_GFP (#61423) by replacing the GFP with a TagBFP gene. SunTag-VP64 plasmids are the Addgene plasmids #60903 and #60904 described above. Primers are listed in Supplementary Table 1.

Target sites for dCas9 were designed through the IDT online design tool (https://www.idtdna.com/SciTools). For cloning target sequence into the corresponding guide RNA vector, protospacer sequence of 20 bp (Supplementary Table 2) was ordered as a pair of complementary oligos with 4 additional nucleotides ACCG- and AAAC-at the 5’ end of the sense and antisense oligonucleotides, respectively. Complementary oligos were annealed by heating at 95°C for 5 min and subsequent cooling to 22°C at a rate of -0.1°C/s. The annealed oligos were then ligated to the BbsI cut site of the vector.

### Lentivirus production and transduction

One day prior to transfection, HEK293T cells were seeded at a density of 1.2×10^6^ cells/well in a 6-well plate in 2 ml Opti-MEM (Invitrogen) containing 2mM GlutaMAX, 1mM Sodium Pyruvate and 5% FCS. Transfection of HEK293T was performed using Lipofectamine 3000 (Invitrogen) as per the manufacturers’ instructions. Cells were co-transfected with packaging plasmids (pCMV-VSV-g and psPAX2) at 0.17 pmol each and around 0.23 pmol transfer construct to make up a final mass of 3.3 µg. Virus was harvested 24- and 52-hour post-transfection. Transduction was performed in a 12-well plate, with 500,000 cells resuspended in 1 ml viral supernatant supplemented with 8 µg/ml polybrene (Millipore). Cells were spun at 2500 rpm at 32°C for 90 minutes. Stable transfectants were enriched by FACS and assayed at the indicated time point, or subjected to further transduction if required.

### Flow cytometry and fluorescence-activated cell sorting (FACS)

For CD4 promoter targeting studies, cells were assayed at the indicated time point post gRNA transduction. Cells were stained with CD4-PE (clone #GK1.5, in-house) and analysed with BD FACSymphony A3. For *Dreg1, Hba-x* and *Hbb-y* studies, cells were sorted on BD FACSAria Fusion or FACSAria III 7 days post gRNA transduction. SAM cells were sorted as mCherry+ BFP+ TagRFP657+ population. SunTag-VP64 or SunTag-TET1 cells were sorted as BFP+ GFP+ TagRFP657+ population. TETact v1-v3 cells were sorted as BFP+ GFP+ mCherry+ TagRFP657+ population.

### Bisulphite sequencing

Genomic DNA was extracted from around 700,000 cells using DNeasy Blood & Tissue kit (Qiagen). 200 ng of gDNA was then subjected to bisulphite conversion and subsequent clean-up using EpiMark Bisulfite Conversion Kit (NEB) as per manufacturers’ instruction. Bisulphite PCR primers for CD4 promoter was designed via Bisulfite Primer Seeker (Zymo) and sequences are listed in Supplementary Table 3. Bisulphite PCR was performed using Phusion U Hot Start DNA polymerase (ThermoFisher) with resultant amplicon gel purified and cloned into pJET1.2 blunt vector (ThermoFisher) of the CloneJET PCR cloning kit (ThermoFisher). Ten clones from each group were analysed via Sanger sequencing.

### Quantitative reverse transcription PCR (RT-qPCR)

RNA was extracted using NucleoSpin RNA Plus (Macherey-Nagel) with gDNA removal. One step RT-qPCR was performed using 20 ng RNA with iTaq Universal probe supermix (Bio-Rad) for Dreg1, or iTaq Universal Sybr Green supermix (Bio-Rad) for CD3ε, Hba-x and Hbb-y, with β-actin as the endogenous reference. Gene expression was normalised to the endogenous control as ΔCT and relative expression evaluated as 2^-ΔCT^. Primers and probes are listed in Supplementary Table 4.

## Author contributions

W.F.C. designed and conducted experiments, C.R.K. T.M.J. provided intellectual input, W.F.C. R.S.A. conceived the study and wrote the manuscript.

## Supplementary Figures

**Supplementary Figure 1.**
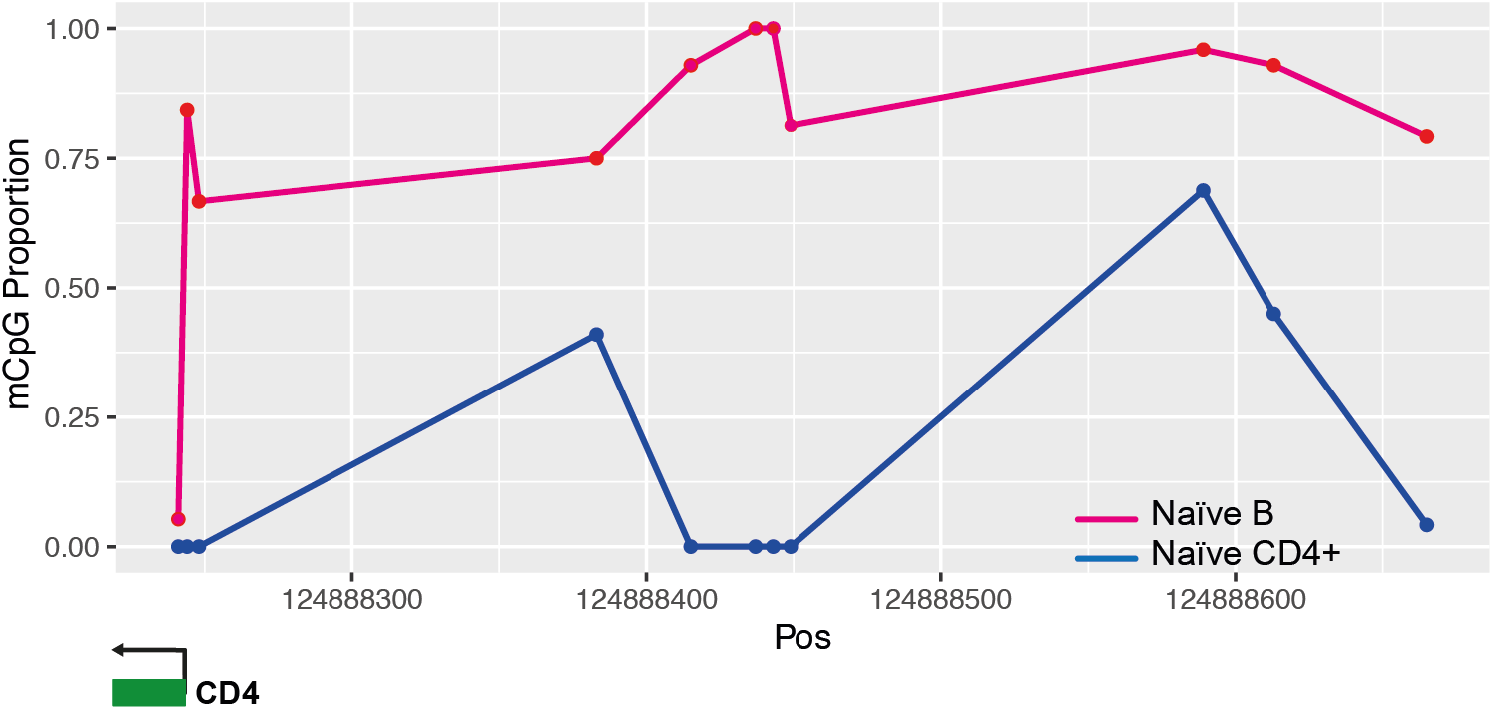
DNA methylation profiles of naïve B and CD4+ T cells at the *CD4* promoter, plotted as population proportion of methylated cytosine in each CpG dinucleotide motif.

**Supplementary Figure 2.**
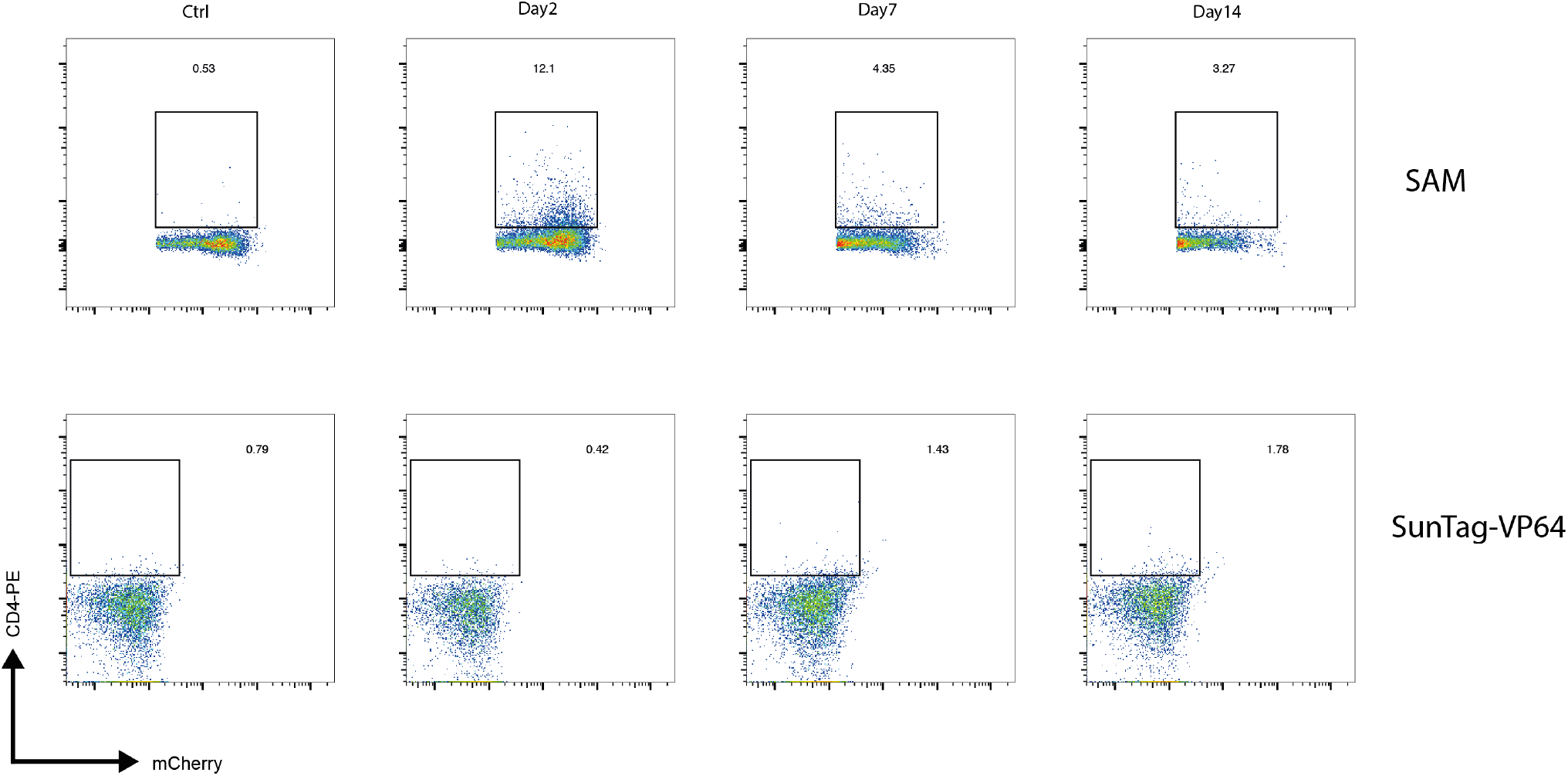
Representative flow cytometry plots showing CD4 surface expression in A20 with SAM or SunTag-VP64 constructs and transduced with CD4-targeting sgRNA on the indicated day post-sgRNA-transduction.

**Supplementary Figure 3.**
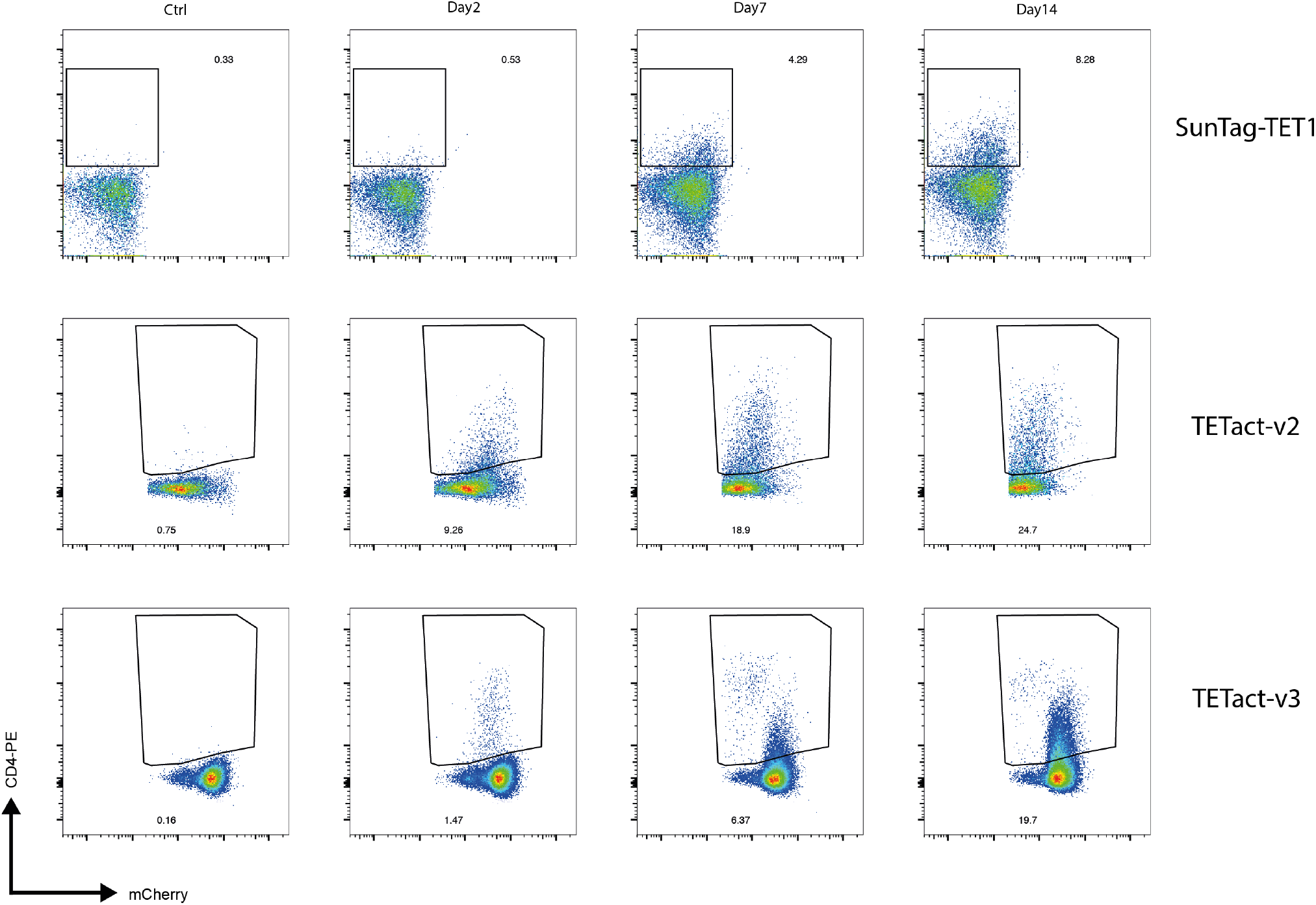
Representative flow cytometry plots showing CD4 surface expression in A20 with SunTag-TET1, TETact-v2 or -v3 constructs and transduced with CD4-targeting sgRNA on the indicated day post-sgRNA-transduction.

**Supplementary Figure 4.**
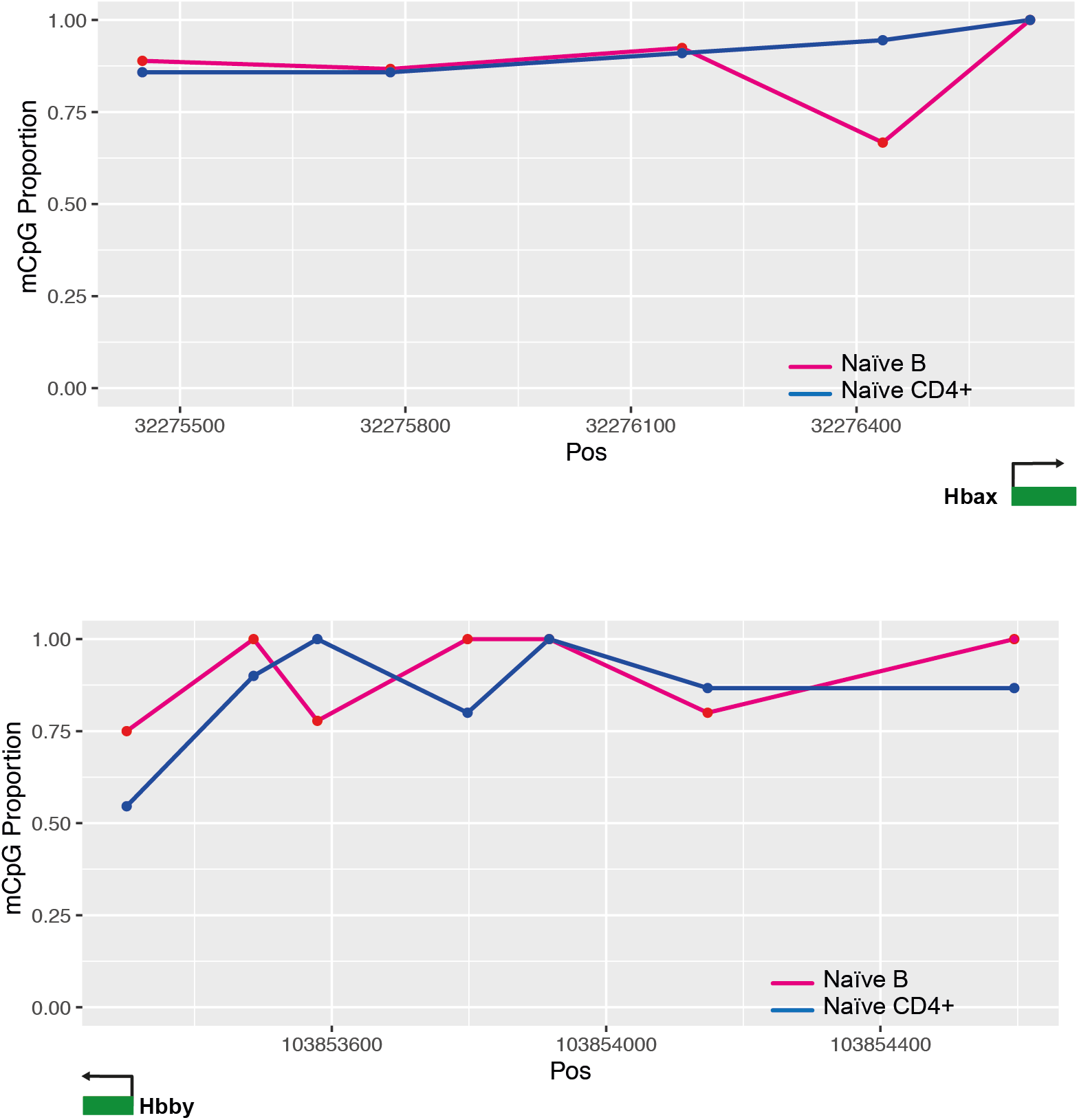
DNA methylation profiles of naïve B and CD4+ T cells at the *Hba-x* (top) and *Hbb-y* (bottom) promoter, plotted as population proportion of methylated cytosine in each CpG dinucleotide motif.

**Supplementary Table 1:**
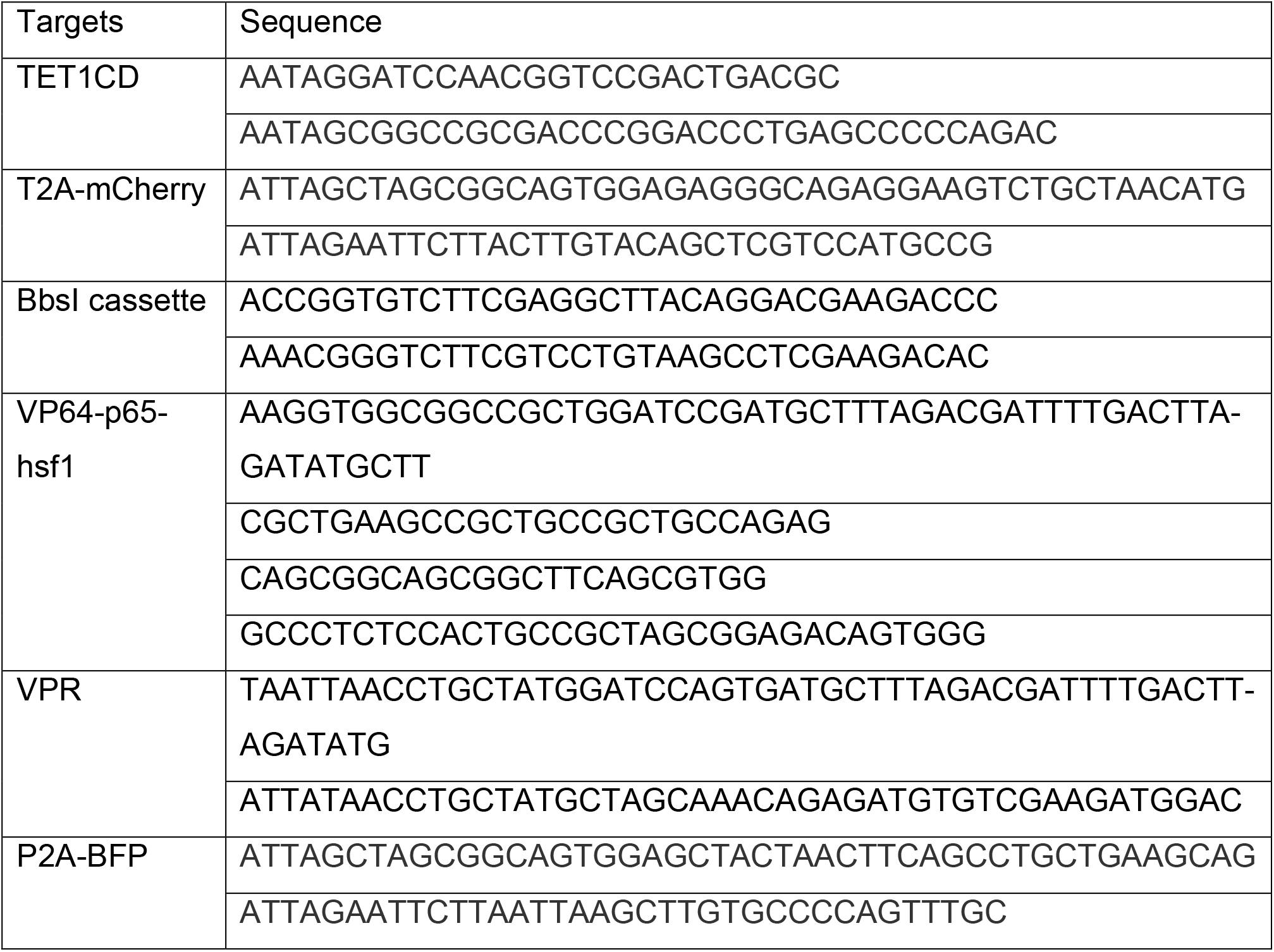
Primers for plasmid construction

**Supplementary Table 2:**
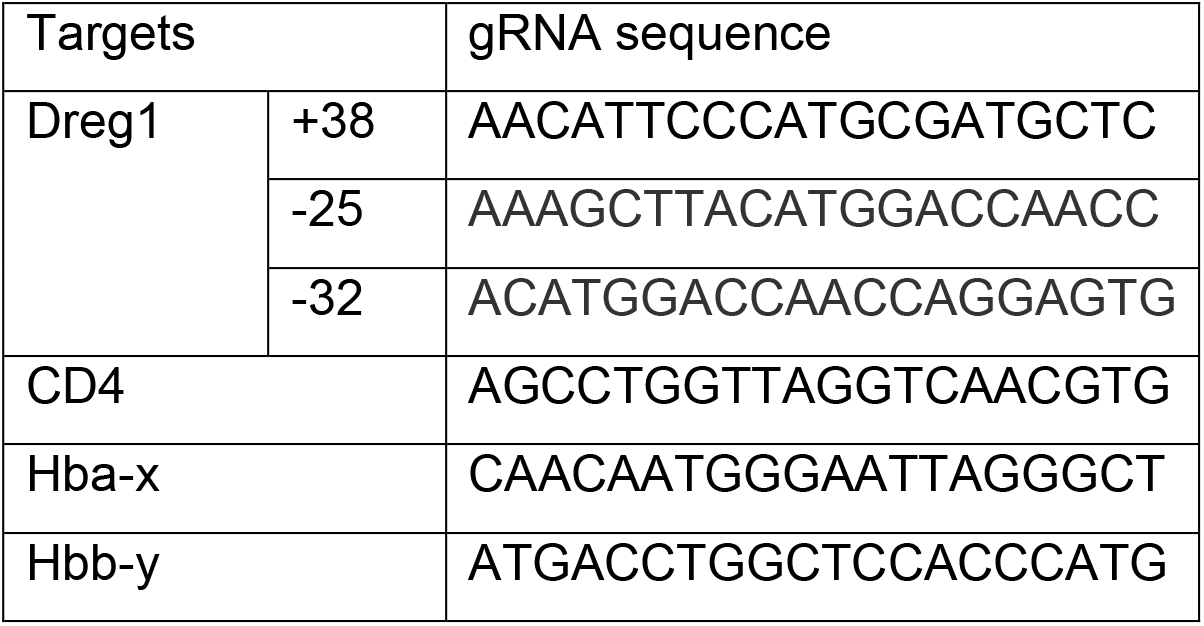
gRNA target sequences

**Supplementary Table 3:**
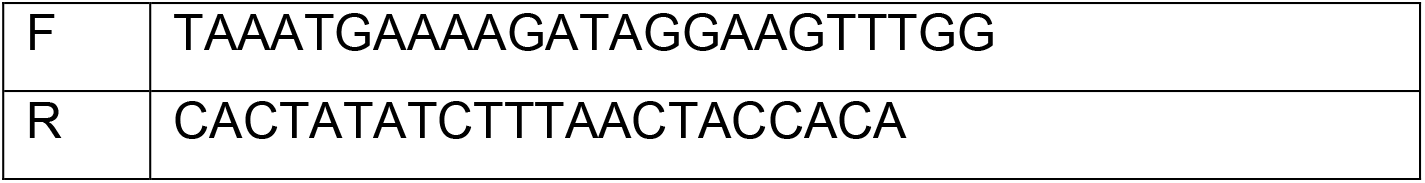
Bisulphite sequencing PCR primers

**Supplementary Table 4:**
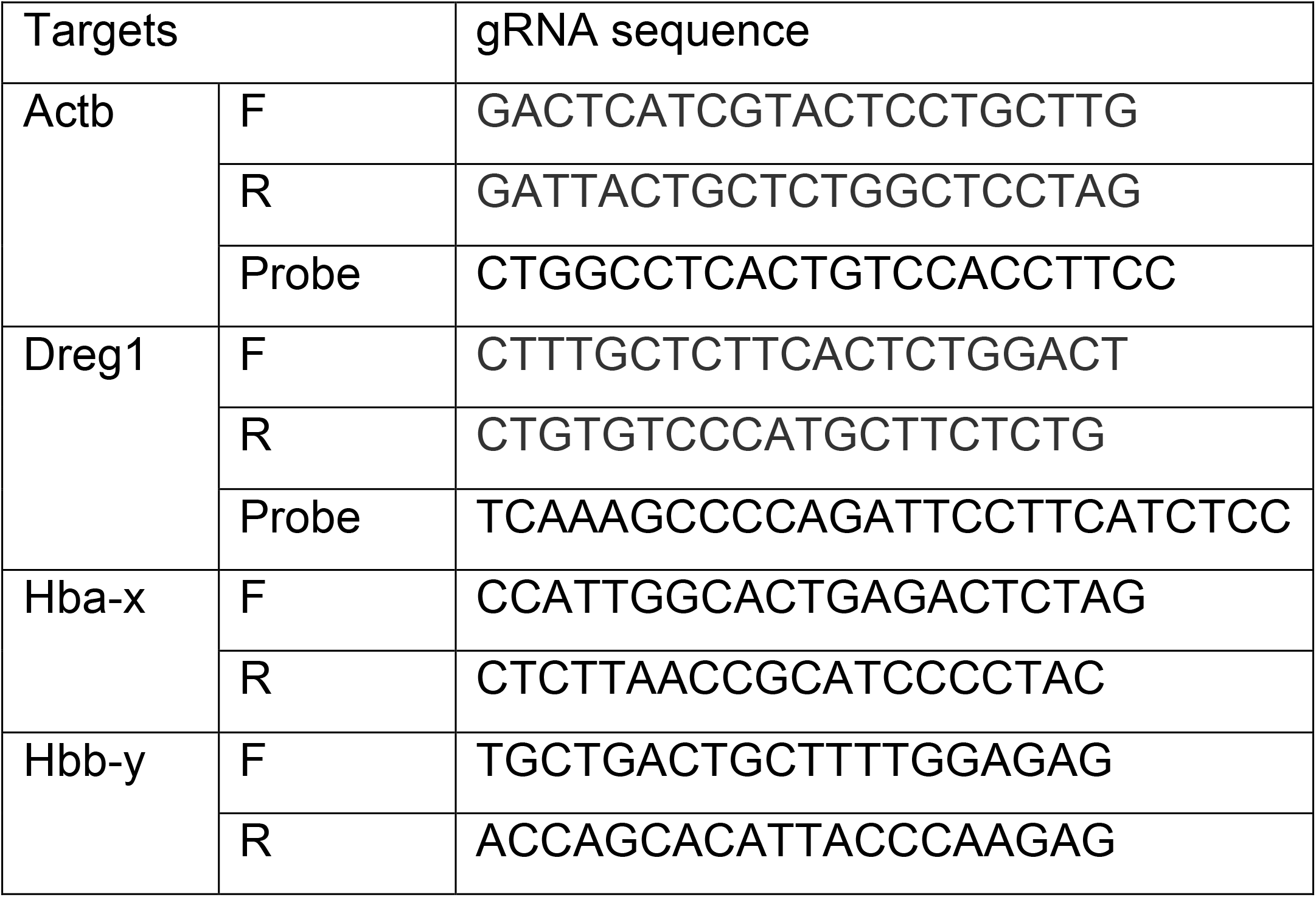
qRT-PCR primers used in study

